# Loss of the high-affinity vacuolar Ca^2+^ pump Pmc1p confers echinocandin tolerance in *Candida albicans* through enhancing calcineurin-based responses

**DOI:** 10.64898/2026.08.26.747206

**Authors:** Justin B. Gregor, Ravinder Kumar, Christian DeJarnette, Glen E. Palmer

**Author notes:** Corresponding author. Mailing Address: University of Tennessee Health Science Center, College of Pharmacy, Department of Clinical Pharmacy and Translational Science, 881 Madison Avenue, Memphis, Tennessee, 38163.

## Abstract

Signaling through the calcium-activated calcineurin phosphatase promotes fungal survival of stressful conditions including those imposed by antifungal medications. Calcium is an essential secondary messenger that regulates diverse cellular processes in eukaryotes; however, it is also profoundly toxic and cytoplasmic concentrations must be tightly controlled. In fungi, the vacuole serves as a major calcium reservoir with the H^+^-exchanger Vcx1p and P-type ATPase Pmc1p sequestering intracellular calcium. Upon stimulation, these stores can be released through the Yvc1p ion channel to create transient cytoplasmic pulses that activate calcium-dependent responses. Despite the importance of calcineurin signaling in sustaining fungal viability upon antifungal insult, the contribution of many other proteins responsible for intracellular calcium homeostasis during antifungal exposure remains poorly understood. Here, we investigated whether Yvc1p, Vcx1p, and Pmc1p influence *Candida albicans* capacity to endure exposure to the first-line echinocandin antifungals. Our results demonstrate that loss of Pmc1p function confers high-levels of tolerance, with *pmc1Δ/Δ* mutant cells sustaining less damage, surviving and capable of proliferation in the presence of supra-MIC concentrations of the echinocandins. Moreover, this phenotype is dependent upon elevated signaling through the calcineurin pathway. Finally, we demonstrate that at least a subset of drugs previously identified as echinocandin antagonists activate calcineurin signaling in a Pmc1-dependent manner to drive echinocandin tolerance. These findings reveal how genetic or pharmacological modulation of calcium homeostasis can have profound consequences on the outcome of *C. albicans* – echinocandin interaction.

## Introduction

The mortality rates for fungal infections that disseminate through the bloodstream are extremely high, exceeding 50% with some pathogenic species (1–3). Several *Candida* species are a leading cause of these invasive fungal infections (IFI’s), as well as common and often recurring mucosal diseases such as oral and vaginal thrush. The modest efficacy of all three antifungal classes used to treat systemic mycoses is a major challenge in their clinical management. For example, approximately a third of patients with disseminated candidiasis are non-responsive to treatment with a front-line echinocandin therapy (4–7), and outcomes are even worse for those treated with an azole antifungal (7). While overt antifungal resistance – defined as a substantial increase in the minimal inhibitory concentration (MIC) above a predetermined clinical breakpoint - contribute to poor patient outcomes (8), ≤ 2% of *Candida albicans* (the most prevalent and virulent *Candida* species) - isolates are echinocandin resistant (1, 8–10). Thus, the majority of treatment failures involve isolates deemed sensitive according to the standard antifungal susceptibility testing protocol.

Although the mechanisms that confer outright echinocandin resistance in *Candida* species (point mutations within the target enzyme that diminish its affinity for these drugs) are well established (8, 11), the mechanisms that promote fungal *survival* following echinocandin exposure are less well characterized. The impact of such ‘tolerance’ mechanisms upon the clinical efficacy of echinocandin therapy, are also unknown. Signaling through the calcium-responsive calcineurin phosphatase has been shown to promote *C. albicans* survival following exposure to both azole and echinocandin antifungals (12–14). However, the role of many other cellular components that facilitate calcium signals in *C. albicans* response to antifungal exposure remains unknown. While calcium is essential for life, excess cytoplasmic Ca^2+^ is acutely toxic and must be quickly removed to avoid cell death (15–17). In contrast to mammals, which extrude most Ca^2+^ out of the cell across the plasma membrane (15), fungi sequester large quantities of excess Ca^2+^ within a specialized intracellular compartment - the fungal vacuole (18, 19). In the yeast *Saccharomyces*, various stimuli including cell wall and antifungal induced stress trigger the release of vacuolar calcium into the cytoplasm through the Yvc1p ion channel, to activate calcium-dependent responses (20, 21). Both low-affinity proton driven (Vcx1p) and high-affinity ATP-driven transporters (Pmc1p) then pump excess Ca^2+^ from the cytoplasm back into the vacuole lumen (20, 22). The goal of this study was to examine how these proteins - which play a central role in the mobilization of intravacuolar calcium stores, influence fungal survival following echinocandin exposure. This in turn can potentially help identify opportunities and define strategies to enhance the therapeutic efficacy of these drugs.

## Results

### Deletion of *PMC1* confers echinocandin tolerance upon *Candida albicans*

Given the central role of calcium-based signaling in promoting fungal survival following antifungal exposure, we examined how dysregulation of vacuolar homeostatic mechanisms affects the antifungal susceptibility of *C. albicans*. Three gene deletion mutants lacking either *VCX1* – encoding the high-capacity low-affinity H^+^-driven transporter that moves Ca^2+^ from the cytoplasm to vacuole lumen; *PMC1* – encoding an ATP-driven low-capacity high-affinity Ca^2+^ transporter; or *YVC1* – encoding a channel that releases Ca^2+^ sequestered within the vacuole into the cytoplasm. According to the CLSI broth microdilution assay, the caspofungin susceptibility of both *vcx1Δ/Δ* and *yvc1Δ/Δ* mutants is indistinguishable from wild-type (figure 1A). However, the MIC of caspofungin was approximately 4-fold-higher for the *pmc1Δ/Δ* mutant (figure 1A). Even more striking, was the robust residual growth of the *pmc1Δ/Δ* mutant observed at supra-MIC concentrations of caspofungin, which was especially prominent after 48 and 72-hours incubation. A similar phenotype was observed with both anidulafungin and micafungin (figure 1B and C).

**Figure 1.**
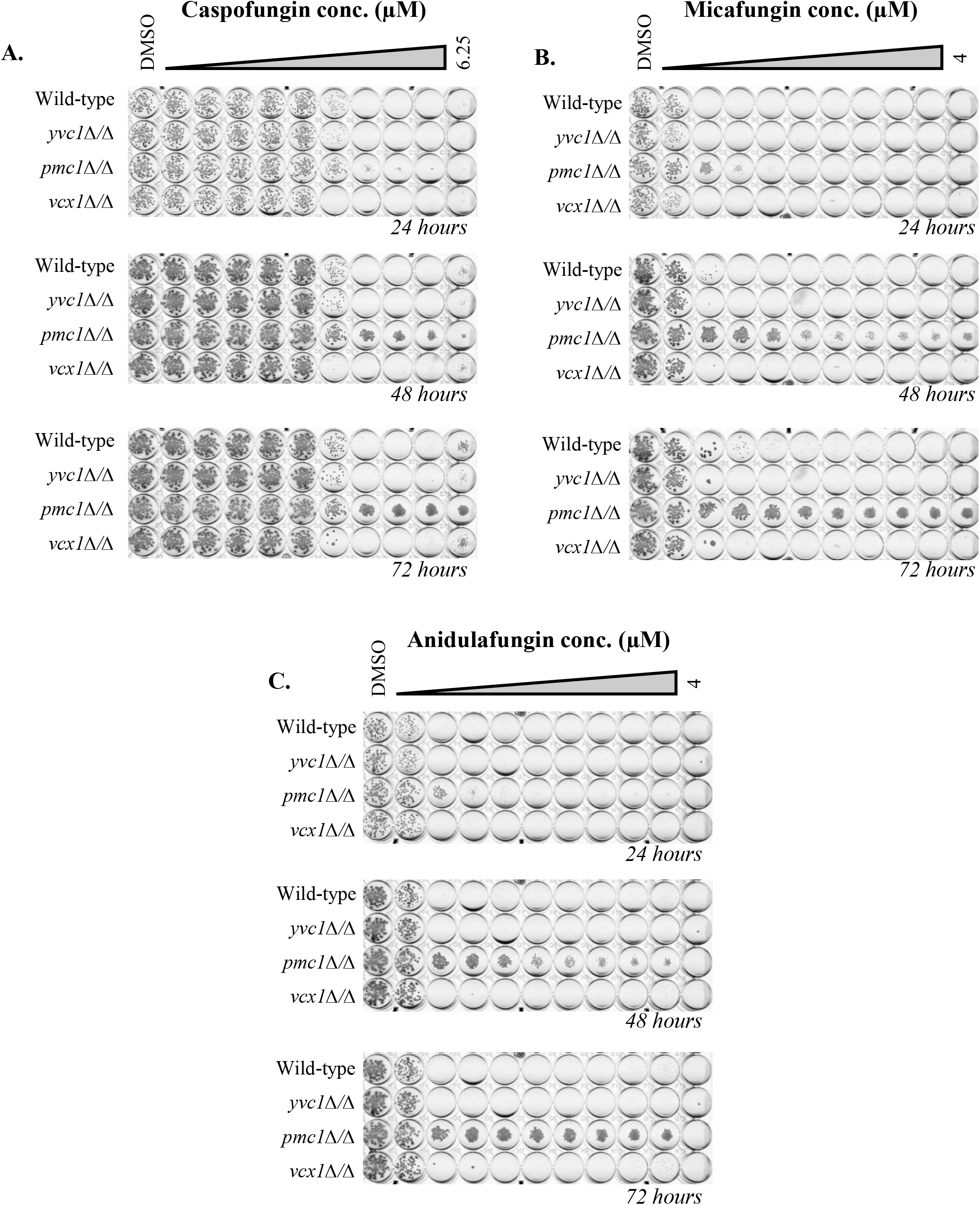
*Candida albicans pmc1Δ/Δ* mutant is echinocandin tolerant. *C. albicans* wild-type, *yvc1Δ/Δ*, *pmc1Δ/Δ* and *vcx1Δ/Δ* strains were seeded into RPMI medium and dispensed into 96-well plates containing a 2-fold dilution series of indicated echinocandin drug, or 0.5% DMSO (vehicle control). Plates were imaged after 24, 48 and 72-hours of incubation at 35°C.

To further characterize the phenotype, growth kinetic assays were performed in the presence of various concentrations of caspofungin. Growth of the wild-type strain was completely halted in the presence of 0.195 or 1.56 µM, but significant growth was observed at 12.5 µM (figure 2), consistent with the previously described ‘paradoxical effect’ (23). In contrast, substantive growth of the *pmc1Δ/Δ* mutant occurred at all caspofungin concentrations. However, the maximum growth rate and cell density attained (OD_600nm_) was less than in the absence of the antifungal (figure 2), indicating the *pmc1Δ/Δ* mutant was not insensitive to caspofungin, but seemingly adept at surviving exposure and capable of continued proliferation. High levels of caspofungin tolerance was also observed with the E-test strip MIC assay (supplemental figure 1).

**Figure 2.**
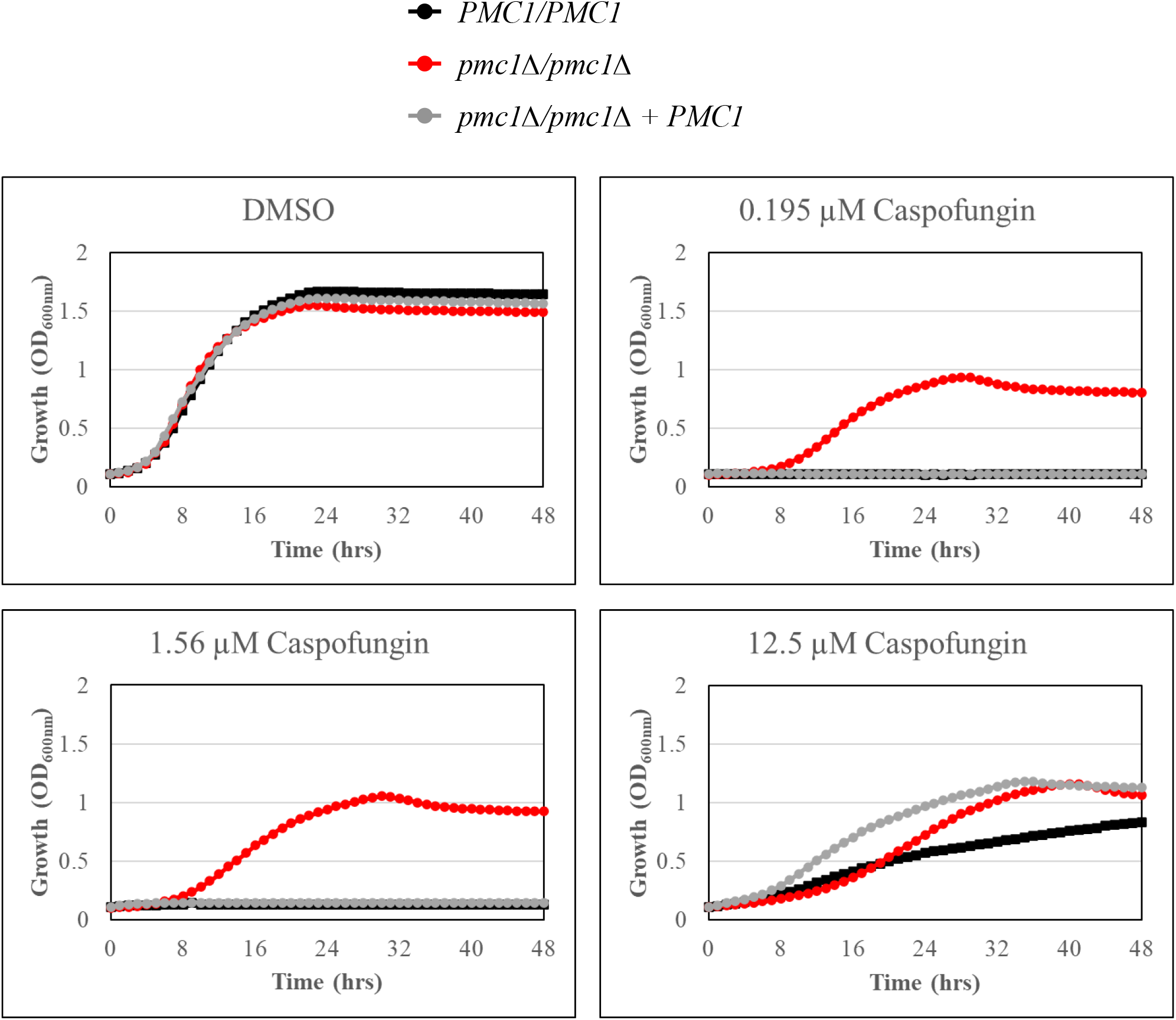
*Candida albicans pmc1Δ/Δ* mutant is able maintain limited growth in the presence of supra-MIC concentrations of caspofungin. *C. albicans* wild-type, *pmc1Δ/Δ* and *pmc1Δ/Δ* + *PMC1* strains were seeded into RPMI medium supplemented with the indicated concentrations of caspofungin, or 0.5% DMSO (vehicle control), incubated at 35°C, and growth measured as OD_600nm_ at 30-minute intervals.

Finally, the mutant exhibited enhanced caspofungin tolerance in YPD (yeast extract peptone) broth, minimal YNB (yeast nitrogen base) medium, and complete synthetic medium (CSM) (figure 3A-C). This indicates that the echinocandin tolerance of the *pmc1Δ/Δ* mutant is a robust phenotype that is not inherently-dependent upon pH or nutrient content of the growth medium.

**Figure 3.**
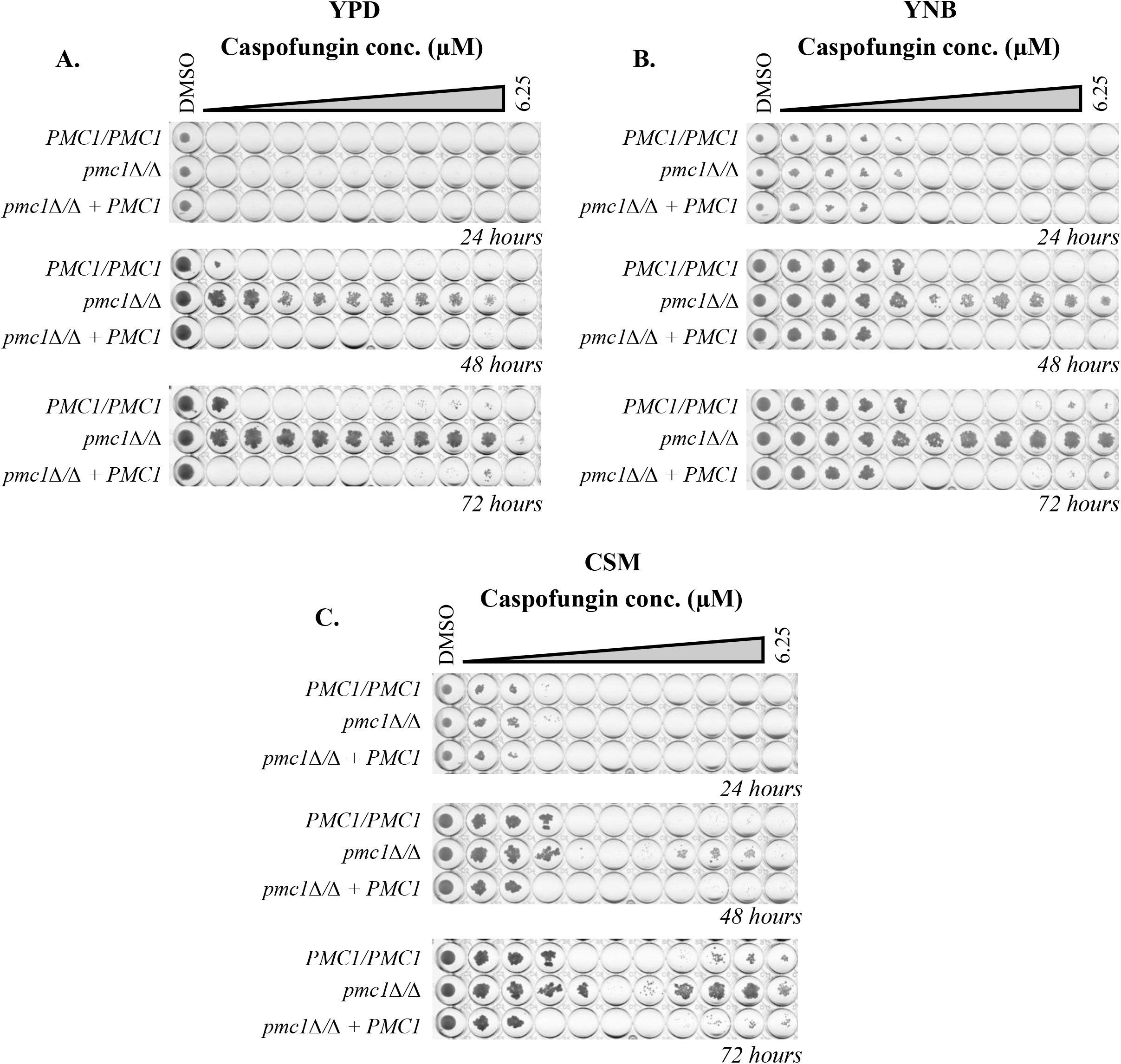
The caspofungin tolerance of *Candida albicans pmc1Δ/Δ* mutant is not a medium specific phenotype. *C. albicans* wild-type, *pmc1Δ/Δ* and *pmc1Δ/Δ* + *PMC1* strains were seeded into (A) YPD, (B) YNB, or (C) CSM media and dispensed into 96-well plates containing a 2-fold dilution series of indicated echinocandin drug, or 0.5% DMSO (vehicle control). Plates were imaged after 24, 48 and 72-hours of incubation at 35°C.

To confirm the relationship between Pmc1p function and echinocandin sensitivity, we constructed *pmc1Δ/Δ* mutants in three clinical isolates, including the reference strain SC5314 (24), ATCC10231 and JS7. The previously reported hyphal growth defects, SDS and calcium sensitive phenotypes (25) were confirmed in all three strain backgrounds (supplemental figure 2). While all three *pmc1Δ/Δ* mutants exhibited reduced caspofungin sensitivity, notable differences were observed in the tolerance phenotype observed. Specifically, the SC5314 and ATCC10231 derived *pmc1Δ/Δ* mutants resembled the lab derived mutant, with elevated residual growth at supra-MIC concentrations of caspofungin, which tapered off at higher concentrations (figure 4). In contrast, the JS7 derived *pmc1Δ/Δ* mutant exhibited what appeared to be an exaggerated paradoxical growth phenotype.

**Figure 4.**
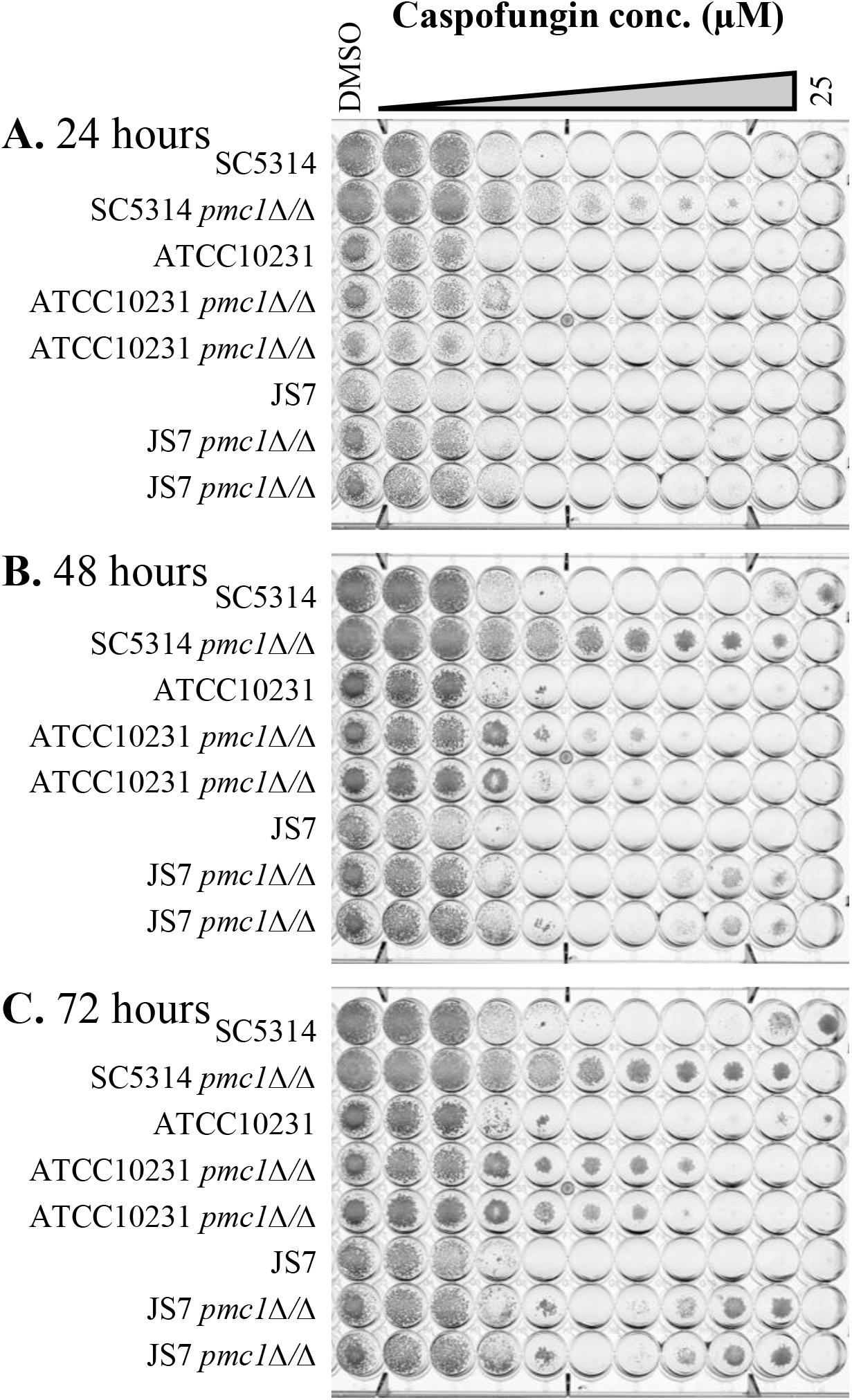
Loss of *PMC1* produces varied caspofungin tolerance phenotypes in different *Candida albicans* isolates. *C. albicans* clinical isolates SC5314, ATCC10231 and JS7, as well as their *pmc1Δ/Δ* derived mutants were seeded into RPMI medium (2% dextrose) and dispensed into 96-well plates containing a 2-fold dilution series of caspofungin, or 0.5% DMSO (vehicle control). Plates were imaged after 24 (A), 48 (B) and 72-hours (C) of incubation at 35°C.

### *Candida albicans pmc1Δ/Δ* mutant cells sustains less damage and survive echinocandin exposure

The viability of mutant and wild-type cells was next compared as colony forming units (CFU) following antifungal exposure. The CFU count for wild-type declined by approximately 1.5 log_10_ units after 4-hours exposure to 4X MIC caspofungin, with a smaller effect observed at 16X MIC, and no significant reduction observed at 64X MIC, again indicating a robust paradoxical effect. In contrast, the viability of the *pmc1Δ/Δ* mutant did not change dramatically over the 4-hours at any concentration of caspofungin (figure 5A). However, the propensity of echinocandin treated cells to aggregate in liquid culture raised concerns that CFUs may not accurately reflect true cell viability. Thus, a second experiment was performed in which untreated cells were plated directly onto agar containing caspofungin. This revealed that essentially all *pmc1Δ/Δ* mutant cells survived and were able to form colonies (95 ± 11%), whereas wild-type yielded few viable colonies (3 ± 1%) (figure 5B). These data indicated that the *pmc1Δ/Δ* mutant exhibited outright caspofungin tolerance with all cells retaining viability, rather than hetero-resistance where a small sub-population of cells survive the antifungal insult.

**Figure 5.**
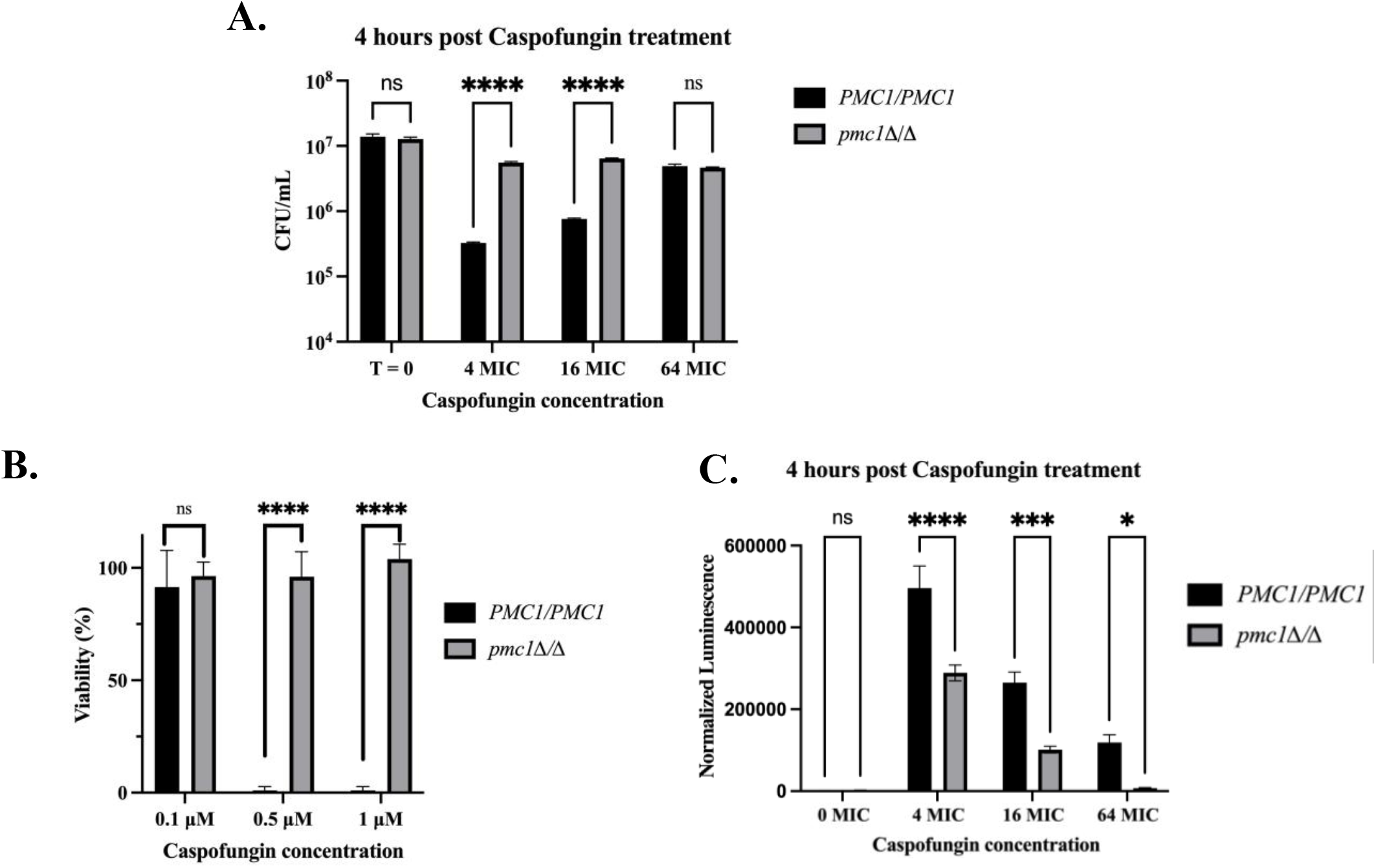
Loss of *PMC1* increases *Candida albicans* resilience to caspofungin induced cell injury. (A) *C. albicans* wild-type and *pmc1Δ/Δ* strains were resuspended in RPMI medium supplemented with 4, 16 or 64X MIC of caspofungin and cell viability quantified as colony forming units (CFU) immediately (t = 0), or after 4-hours incubation at 35°C (t = 4 hours). (B) Wild-type and *pmc1Δ/Δ C. albicans* strains were plated onto RPMI agar supplemented with 0.1, 0.5, or 1 µM caspofungin. Colonies were counted after 48 hours at 35°C and viability colony formation in the presence of each antifungal concentration expressed as a percentage of colony formation in the absence of caspofungin (0.5% DMSO - vehicle control). (C) Wild-type and *pmc1Δ/Δ* strains expressing Nluc were resuspended into RPMI medium supplemented with 0 (0.5% DMSO), 4, 16 or 64X MIC of caspofungin and levels of Nluc activity released into cell-free culture supernatant measured after 4-hours incubation at 35°C. Data presented are the mean of 3 independently conducted experiments and normalized to OD_600_ reads. Error bars represent standard deviation. P-values were calculated from two-way ANOVAs with multiple comparisons and are represented as follows: ns, P > 0.05; *, P < 0.05. ns, ****, P<0.0001.

Levels of cellular damage sustained by populations of cells was next compared following antifungal exposure. *C. albicans* strains expressing a small cytoplasmic luciferase (NanoLuc™, Promega) (26) were used, and levels of enzyme released into the culture supernatant measured after 4-hours exposure to either 4X, 16X or 64X MIC of caspofungin. Exposure of wild-type to 4X MIC of caspofungin increased NanoLuc release by approximately 100-fold compared to mock-treated controls (figure 5C), indicating substantial levels of cell injury and/or lysis. Interestingly, progressively lower levels of Nluc release occurred at higher concentrations of the antifungal, indicating that the paradoxical effect manifests as reduced levels of cell damage as well as residual growth of *C. albicans*. At all concentrations, the *pmc1Δ/Δ* mutant released much less Nluc, indicating it sustained significantly less injury than wild-type. Collectively, these data establish that the *pmc1Δ/Δ* mutant sustains less damage than wild-type, is not prone to the cidal effects of, and is capable of continued growth in the presence of supra-MIC concentrations of the echinocandins.

### Echinocandin tolerance of the *pmc1Δ/Δ* mutant is dependent upon elevated calcineurin signaling

Supplementing the medium with calcium, or sequestering it with EGTA, had little impact upon the echinocandin tolerance of the *pmc1Δ/Δ* mutant (figure 6A). However, inclusion of FK-506 or cyclosporin A, both inhibitors of calcineurin signaling, restored caspofungin sensitivity to the *pmc1Δ/Δ* mutant (figure 6B and 6C). To further examine how loss of Pmc1p influences calcineurin signaling, we utilized a GFP-reporter coupled to the 5’ UTR of the calcineurin responsive *RTA2* gene (27). In both YPD and RPMI medium, *P_RTA2_-GFP* reporter activity was essentially undetectable in wild-type, while the addition of 0.1 µM caspofungin steadily induced the reporter ∼5-20-fold (figure 7A and 7B). In contrast, even in the absence of caspofungin, GFP levels in the *pmc1Δ/Δ* background approximate those observed in the wild-type with caspofungin. Addition of caspofungin further elevates GFP expression in the *pmc1Δ/Δ* mutant by ∼5-20-fold (figure 7A and 7B). These data indicate that calcineurin signaling is substantially elevated in the *pmc1Δ/Δ* mutant and required for its echinocandin tolerance.

**Figure 6.**
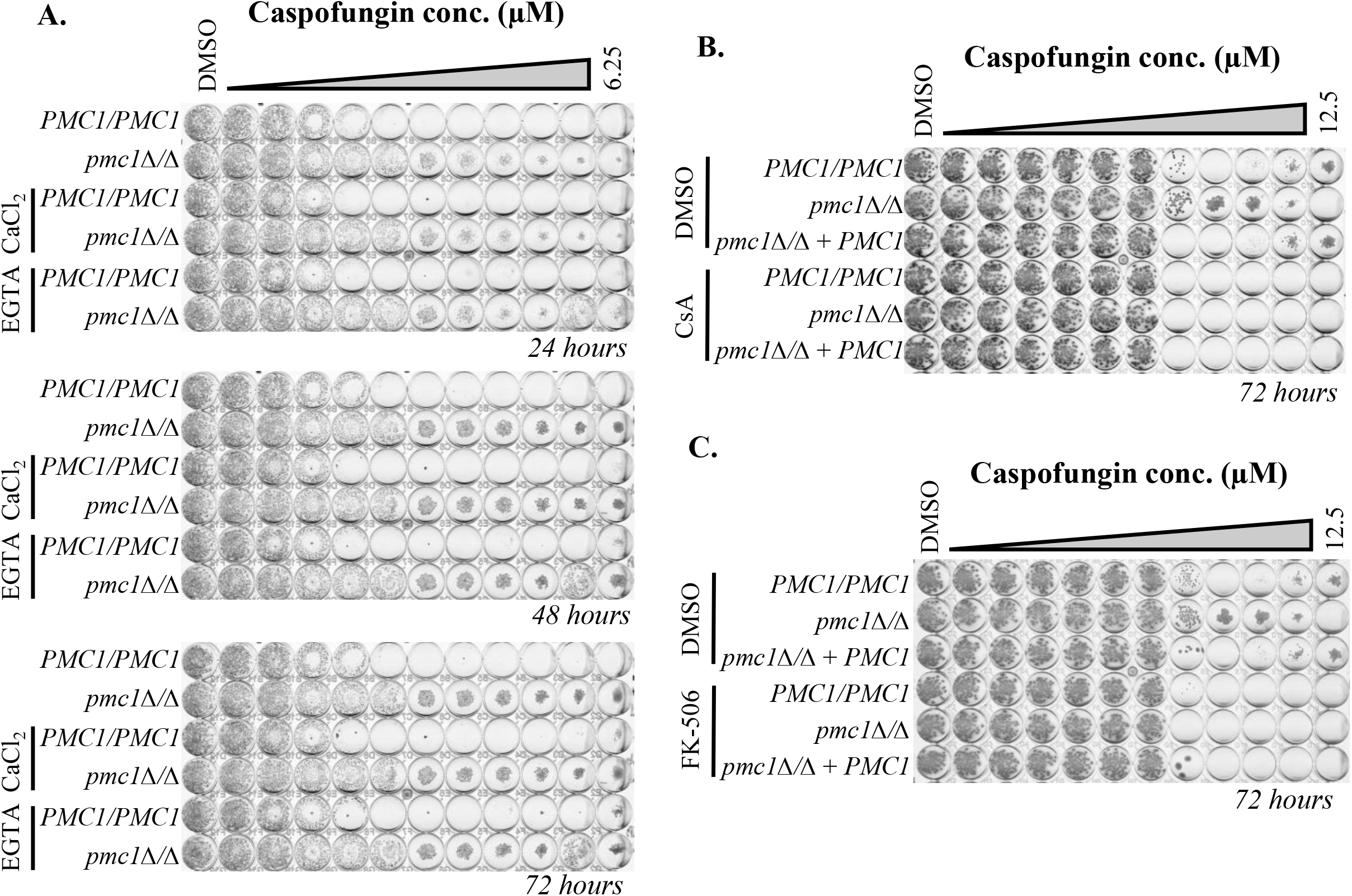
Echinocandin tolerance of the *pmc1Δ/Δ* mutant is dependent upon calcineurin signaling. (A) *C. albicans* wild-type and *pmc1Δ/Δ* strains were seeded into RPMI medium and dispensed into 96-well plates containing a 2-fold dilution series of caspofungin or 0.5% DMSO (vehicle control). Medium was supplemented with water (vehicle control), 200 µM CaCl_2_, or 100 µM EGTA, and plates imaged after 24, 48 and 72-hours of incubation at 35°C. (B and C) Assays were set up as described in panel A, except the medium was supplemented with either 0.5% DMSO or 8 µg/mL cyclosporin A (CsA - panel B), or 1 µg/mL FK-506 (panel C), and plates imaged after 72 hours.

**Figure 7.**
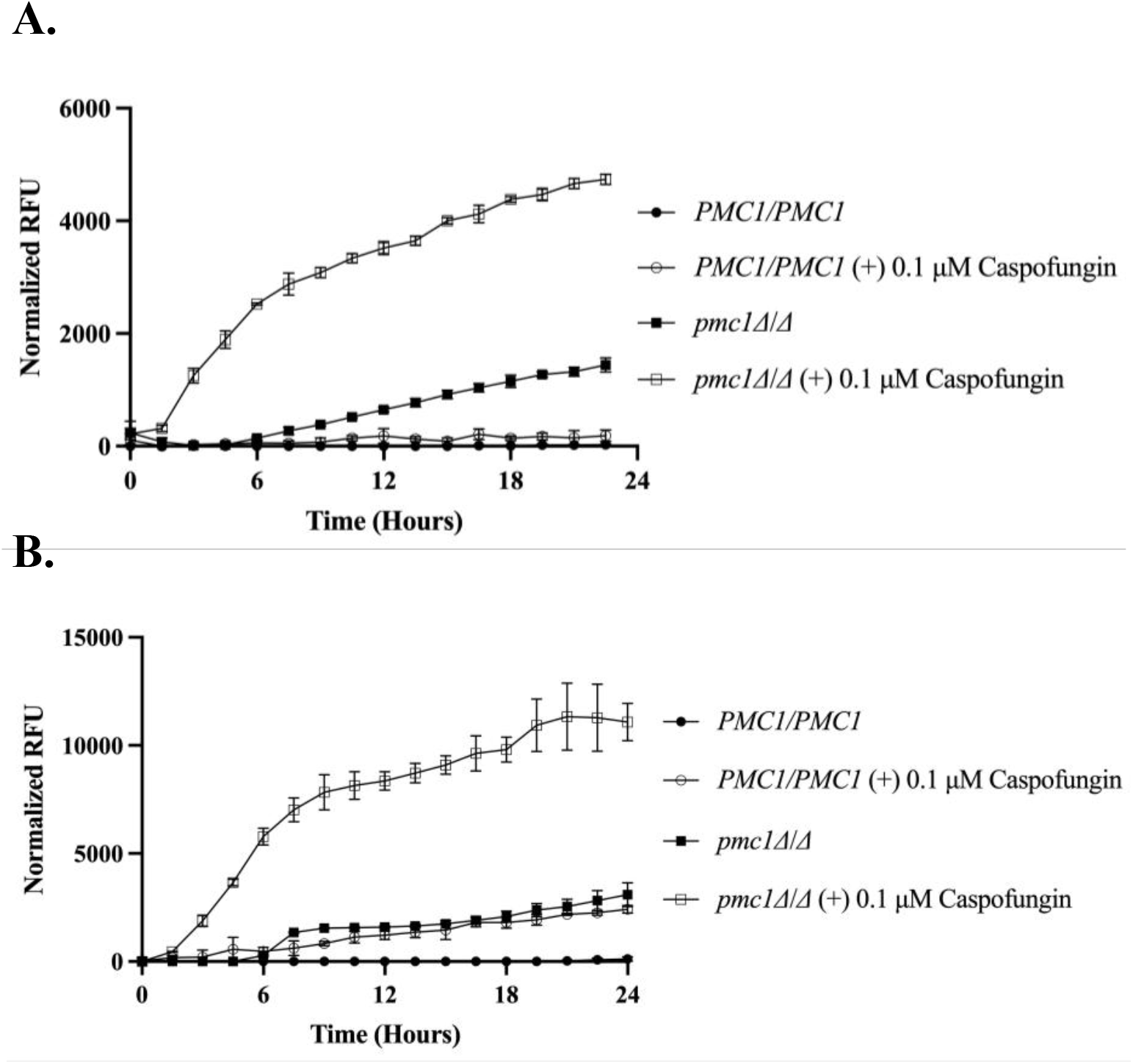
Loss of *PMC1* elevates basal and caspofungin-induced levels of calcineurin signaling. *C. albicans* wild-type and *pmc1Δ/Δ* cells expressing the *P_RTA2_-GFP* reporter were seeded into RPMI (A) or YPD medium (B) supplemented with 0.5% DMSO (vehicle control) or 0.1 µM caspofungin and cells incubated at 35°C. GFP fluorescence intensity and growth (OD_600nm_) were measured at 90-minute intervals. Background fluorescence was subtracted and GFP signal normalized to OD_600nm_. Data presented are the mean of three independently conducted experiments, with error bars indicating standard deviation. Area under the curve was measured and One-way ANOVA with multiple comparisons was performed. All groups are statistically different at P < 0.001.

### Haloperidol and ponatinib activate calcineurin signaling in *Candida albicans* to promote echinocandin tolerance

We previously reported a collection of medications approved for human use that antagonize the antifungal activity of the echinocandins (28). Many of these medications promote substantive levels of residual growth above MIC, resembling the phenotype of the *pmc1Δ/Δ* mutant. To determine if the mechanisms underlying this echinocandin antagonism overlap with the *pmc1Δ/Δ* mutant’s tolerance, we selected a subset of five and determined if they impinged upon calcineurin signaling using the *P_RTA2_-GFP* (27). Of the five drugs tested, the antipsychotic haloperidol and tyrosine kinase inhibitor ponatinib, both induced FK-506-repressible GFP expression by roughly 5-10 fold in a dose dependent manner (figure 8A and C), indicating they stimulate signaling through the calcineurin pathway. Neither drug stimulated *RTA2-GFP* expression above the already elevated levels in the *pmc1Δ/Δ* mutant (figure 8B and D). Collectively, these data indicate that both haloperidol and ponatinib induce calcineurin signaling in a Pmc1p-dependent manner to promote echinocandin tolerance.

**Figure 8.**
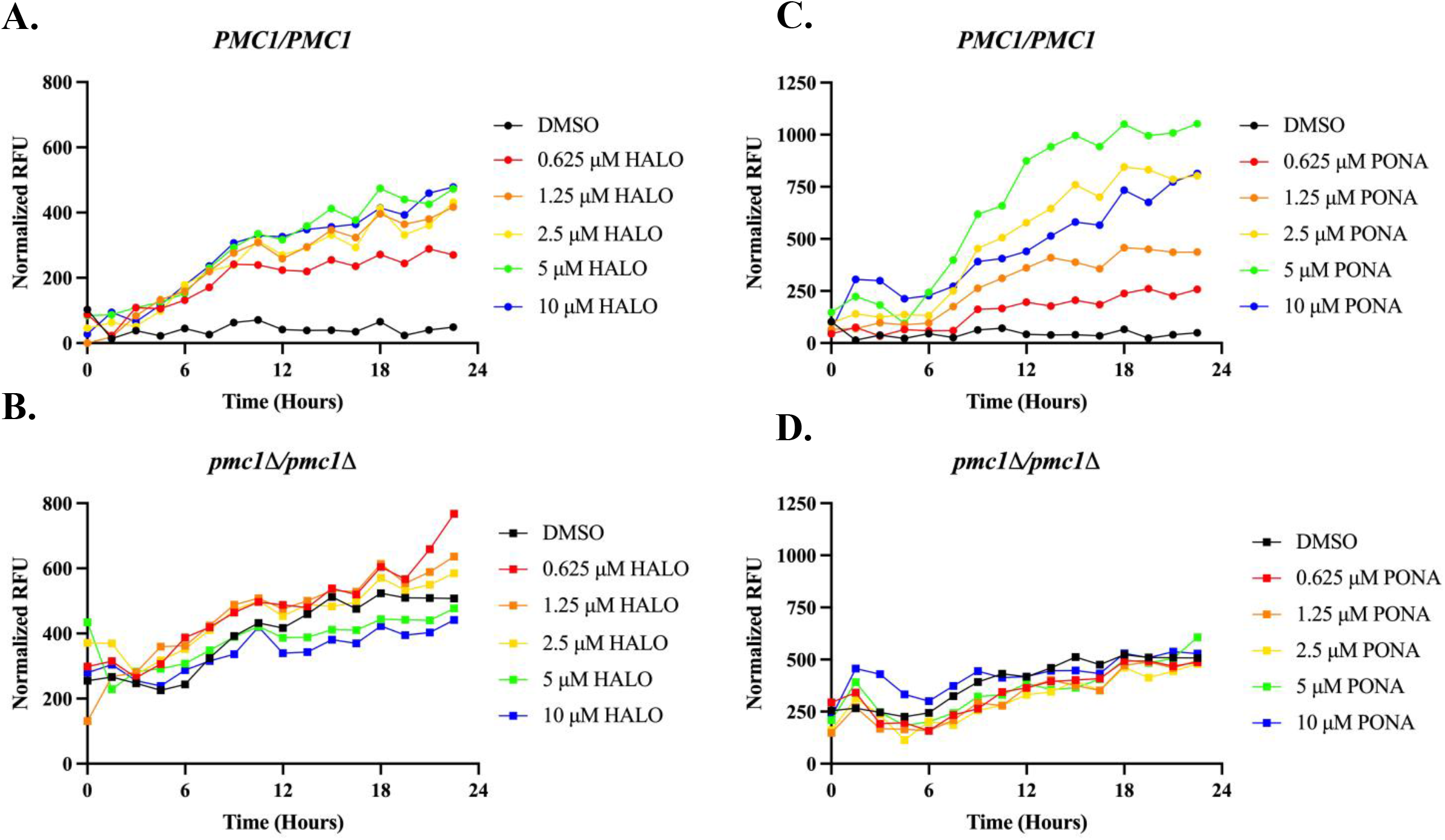
The echinocandin antagonists haloperidol and ponatinib induce calcineurin signaling in a Pmc1p-dependent manner. *C. albicans* wild-type (A and C) and *pmc1Δ/Δ* strains (B and D) expressing *P_RTA2_-GFP* reporter were seeded into RPMI medium (2% dextrose) supplemented with 0.5% DMSO (vehicle control) or indicated concentrations of haloperidol or ponatinib. Data was normalized to OD_600_ reads, and data was set relative to wild-type + indicated concentration of drug to control for background fluorescence. Data presented are the mean of 3 independently conducted experiments.

## Discussion

Signaling through the calcineurin pathway has long been known to promote fungal survival of stressful conditions, including those resulting from antifungal insult. This has raised substantial interest in targeting the calcineurin phosphatase as a strategy to enhance antifungal potency (14, 29–31). The work described herein highlights that modulation of calcium homeostasis, either through natural and heritable variation in gene function, or by pharmacologically active substances that can alter basal and stress-induced levels of calcineurin signaling, and therefore the capacity of fungi to survive antifungal insult.

We have previously shown that the *C. albicans pmc1Δ/Δ* mutant has elevated levels of cytoplasmic calcium (25), presumably due to a diminished capacity to sequester calcium within the vacuole. Our data indicate that essentially all *pmc1Δ/Δ* cells survive echinocandin exposure rather than a sub-population, which precludes classification of this phenotype as heteroresistance. In addition, the *pmc1Δ/Δ* cells are able to grow in the presence of supra-MIC concentrations of the echinocandins, albeit at a rate below drug-free cultures. As such, we have classified the *pmc1Δ/Δ* mutants’ phenotype as echinocandin tolerance. Calcineurin signaling is absolutely required for the mutants echinocandin tolerance. Further, the *RTA2-GFP* reporter data are consistent with the elevated levels of cytoplasmic calcium in the *pmc1Δ/Δ* mutant enhancing both the basal and echinocandin-induced calcineurin signaling that trigger downstream effector responses of sufficient intensity to confer protection from the normally cidal effects of the echinocandins. Interestingly, although calcineurin signaling is known to promote *C. albicans* survival of the fungistatic azole antifungals, the *pmc1Δ/Δ* mutants’ sensitivity to these antifungals was unaffected under the conditions of the CLSI susceptibility assay (Supplemental figure 3A and B) (27). This suggests that quantitative and/or qualitative differences in calcineurin-based responses confer protection against each antifungal drug class.

A key question remaining is whether the echinocandin tolerance of the *pmc1Δ/Δ* mutant, or isolates exhibiting echinocandin tolerance more generally, are of clinical significance. Specifically, whether the tolerance observed *in vitro*, translates to reduced therapeutic efficacy within an infected patient. Our previous work demonstrated that the *pmc1Δ/Δ* mutant has substantially diminished virulence in a mouse model of disseminated candidiasis (25). The mutant also has reduced capacity to form hyphae, which has been associated with tissue damage (32), and exhibits sensitivity to calcium and membrane stress. Thus, it is clear that complete loss of Pmc1p function incurs a significant fitness cost. However, the virulence studies were conducted using immunocompetent mice, and it remains plausible that the mutant is able to cause a productive infection within an immunosuppressed host. Thus, despite a significant fitness trade-off, it is possible loss of Pmc1p function could confer reduced echinocandin sensitivity. Furthermore, complete loss of Pmc1p function may not be necessary to enhance *C. albicans* echinocandin tolerance. Alternatively, compensatory mutations could potentially diminish the fitness costs associated with loss of Pmc1p function, without negating the potential benefits of the tolerance phenotype. Thus, an important role for Pmc1p and related factors that control calcium homeostasis and signaling as determinants of echinocandin therapeutic efficacy cannot be ruled out. With over 30% of invasive *C. albicans* infections failing echinocandin therapy, but less than 2% exhibiting outright resistance (1, 8–10), it is crucial to understand how antifungal tolerance influences clinical outcomes.

It has become increasingly apparent that significant variation in genotype and organization of various signaling networks underly sometimes disparate phenotypes and physiological responses of *C. albicans* clinical isolates (33). This is likely to include variation in the expression level of Pmc1p as well as functional deviation of its allelic variants, its regulatory factors and other cellular components involved in calcium homeostasis. Such variation may impact both the basal and stress-induced levels of cytosolic free-calcium, the duration and temporal regulation of the resulting calcium pulses, as well as the thresholds required for signal and effector activation. Thus, we anticipate natural genetic variation to be a significant source of heterogeneous outcomes of *C. albicans*-echinocandin interaction – specifically the rate and extent of cell killing. Our results are largely consistent with those recently reported by Cunningham and colleagues (34), who found that activation of calcineurin signaling through genetic or pharmacologically induced-stress promoted echinocandin tolerance in *Candida glabrata*. One notable difference from our *C. albicans pmc1Δ/Δ* mutant is that the phenotype they described was a property of a sub-population of the *C. glabrata* cells. Nonetheless, in previous studies we have identified a large collection of drugs approved for human use that induce an echinocandin tolerance-like phenotype (28). The findings of this study suggest that a subset of these medicines activate signaling through the calcineurin pathway in a manner analogous to that observed within the *pmc1Δ/Δ* mutant.

Specifically, haloperidol and ponatinib activated calcineurin signaling. Thus, in addition to genetic or epigenetic variation between fungal isolates, there is significant potential for inter-patient variation in drug-induced calcium-calcineurin-based responses that could potentially effect antifungal responses, and ultimately therapeutic efficacy. Haloperidol is known to stimulate influx of calcium into the cytoplasm of mammalian cells through L-type calcium channels (35, 36), and more recently shown to elevate cytoplasmic calcium in *C. albicans* (37). Such drug-induced activation of calcineurin signaling within invading fungal pathogens is especially concerning given the extensive, complex and patient-specific nature of drug regimens administered to patients at the greatest risk of invasive fungal infections. Curiously, we have observed that some drugs known to target components of the calcium homeostatic and signaling mechanisms within eukaryotic cells enhance echinocandin tolerance (i.e. elevate residual growth above the MIC) and azole resistance (i.e. elevate MIC). For example, the calcium mimetic cinacalcet and the L-type calcium channel niguldipine (Supplemental figure 3). In contrast, other drugs known to perturb calcium homeostasis including benzbromarone and benziodarone, have effects that are specific to the antifungal class (Supplemental figure 3)(38).

Together with emerging evidence of echinocandin-related phenotypes across the *Candida* genera (34, 39), these findings underscore that fungal responses to echinocandins extend beyond the traditional framework of susceptibility and resistance for what has been classed as a “fungicidal” drug. Although currently poorly characterized, tolerance, heteroresistance and paradoxical growth may provide an important means by which *Candida* spp. adapt to drug exposure. A more comprehensive understanding of these phenotypes and the mechanisms that govern them will provide desperately needed insight into the causes of treatment failure and identify opportunities to improve the clinical outcomes of therapeutic intervention.

## Methods

### Growth conditions

*C. albicans* was routinely grown at 30°C in YPD medium (1% yeast extract, 2% peptone, 2% dextrose) supplemented with uridine (50 μg/ml) when necessary. Transformant selection was carried out on minimal YNB medium (6.75 g/liter yeast nitrogen base without amino acids, 2% dextrose, 2% Bacto agar) supplemented with the appropriate auxotrophic requirements, as described previously for *Saccharomyces cerevisiae* (40), or 50 μg/ml uridine.

### Strain construction

All strains used in this study are listed in Table S2. Transformation of *C. albicans* with DNA constructs was performed using the lithium acetate method (41). Gene deletion strains were constructed by the PCR-based approach described by Wilson et al. (42).

### Growth kinetic assays

Growth curve assays were set up in 96-well plates using RPMI-pH 7 (2% glucose). *C. albicans* strains were grown overnight in YPD at 30°C, and the cell density was adjusted to 1 × 10^4^ cells/mL in the appropriate medium for the growth kinetic assays. An aliquot of cell suspension (100 µL) was mixed with an equal volume of twice the desired drug concentration of each antagonist. Cells were then incubated at 35°C inside a BioTek Cytation 5 plate reader with shaking for 48 hours, and OD_600_ was read every 30 min. Data were then analyzed via GraphPad Prism software. These assays were repeated in biological duplicate.

### Antifungal susceptibility assays

Antifungal susceptibility testing was performed using the broth microdilution method as described in Clinical and Laboratory Standards Institute document M27-A3 (43), with minor modifications. Each echinocandin was diluted in DMSO at 100x the final concentration and serially diluted in a round-bottom 96-well plate. *C. albicans* strains were grown overnight in YPD at 30°C, resuspended at 1 × 10^4^ cells/mL in RPMI-pH 7 with 2% glucose, and 100 µL was transferred to wells of a round-bottomed 96-well plate containing concentrated echinocandin solution. The final concentration of DMSO was 0.5% for all treatments, with drug-free control wells having DMSO alone. Plates were incubated without shaking for 72 hours at 35°C, with plates scanned every 24 hours using an EPSON Perfection v.700 Photo scanner. Experiments were performed in biological duplicate.

### Cell survival, plating, and cell damage assays

For cell survival assays, indicated strains were grown overnight in YPD at 30°C. Cells were washed and subcultured at 1 × 10^6^ cells/mL in 10 mL of RPMI-pH 7 (2% glucose) supplemented with 0.5% DMSO (vehicle), 0.4 µM, 1.6 µM, or 6.25 µM caspofungin alone. Cells were incubated at 35°C for 4 hours at 250 rpm plus or minus drug with shaking. After incubation, cells were spun down at 3,000 revolutions per minute (rpm) for 5 min and washed twice with sterile deionized water. Cells were then resuspended in sterile deionized water and serially diluted, and 100 µL of cell solutions was plated onto YPD plates. Plates were then incubated for 48 hours at 30°C, and cell viability was measured by counting colony-forming units (CFUs). For cell damage assays, cells were grown under the same conditions, and a portion of supernatant was collected for measuring luciferase activity. Experiments were repeated in biological triplicate. For cell damage assays, supernatant was collected from the cells grown in the previously listed condition. Luciferase-based assays were performed as previously described (27). Experiments were performed in biological triplicate, and statistical significance was calculated via one-way analysis of variance (ANOVA) test. For caspofungin plating assays, indicated strains were grown overnight in YPD at 30°C. Cells were washed, resuspended in RPMI-pH 7 (2% glucose), and 200 cells were plated directly onto RPMI RPMI-pH 7 (2% glucose) agar containing 0.5% DMSO (vehicle), 0.1 µM, 0.5 µM, or 1 µM caspofungin. Plates were then incubated for 48 hours at 30°C, and cell viability was measured by counting colony-forming units (CFUs).

### Fluorescence Reporter Assays

*C. albicans* strains expressing GFPγ from the *RTA2* promoter were grown overnight in YPD at 30°C. Cells were washed and resuspended in RPMI + 2% glucose to an OD_600nm_ of 0.1 and allowed to grow to mid-log phase. Cells were then back-diluted to an OD_600nm_ of 0.1 and 100 μL of cells were dispensed into a round-bottomed 96-well plate. Cells were treated with either 1 μM caspofungin 0.5% DMSO, or indicated concentrations of Haloperidol/Ponatinib from 0 to 10 μM and incubated at 35°C for 24 h. GFPγ fluorescence intensity was then quantified by using a Cytation 5 plate reader (BioTek Instruments, Inc.) with excitation at 488 nm and emission at 507 nm, each with a 9 nm bandwidth. Growth was determined by measuring the OD_600_ for normalization of the fluorescence signal.

### Hyphal Growth and Spot-dilution assays

Each *C. albicans* strain was grown overnight in YPD at 30°C. The cells were washed in sterile deionized water, the cell density was adjusted to 10^7^ cells/ml, and 1:5 serial dilutions were performed in a 96-well plate. Each cell suspension was then applied to agar plates using a sterile multipronged applicator. Resistance to different stresses was determined on YPD agar containing 0.05% SDS or 500 mM CaCl_2_, with incubation at 30°C for 48 to 96 h. To induce hyphal growth, for each strain, 2.5 µl of a 10^7^ cells/ml cell suspension was spotted on M199 or 10% fetal bovine serum (FBS) agar plates, and incubated for 96 h at 35°C.

## Acknowledgements

This work was funded by the National Institute of Allergy and Infectious Diseases of the National Institutes of Health under award number R01AI152067. The content is solely the responsibility of the authors and does not necessarily represent the official views of the National Institutes of Health.

---

**Supplemental figure 1.**
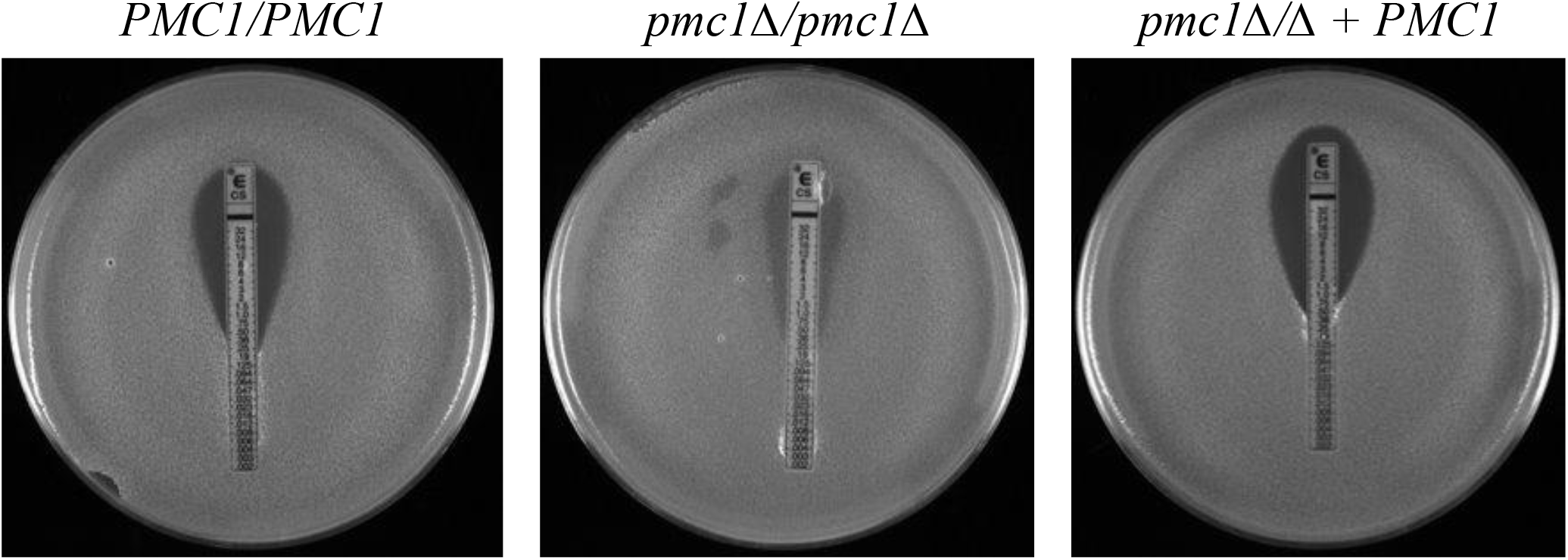
*Candida albicans pmc1Δ/Δ* mutant is echinocandin tolerant in E-test assay. *C. albicans* wild-type, *pmc1Δ/Δ* and *pmc1Δ/Δ* + *PMC1* strains were spread onto RPMI agar plates. E-test strips were placed on agar plates and plates imaged after 24-hours of incubation at 35°C.

**Supplemental figure 2.**
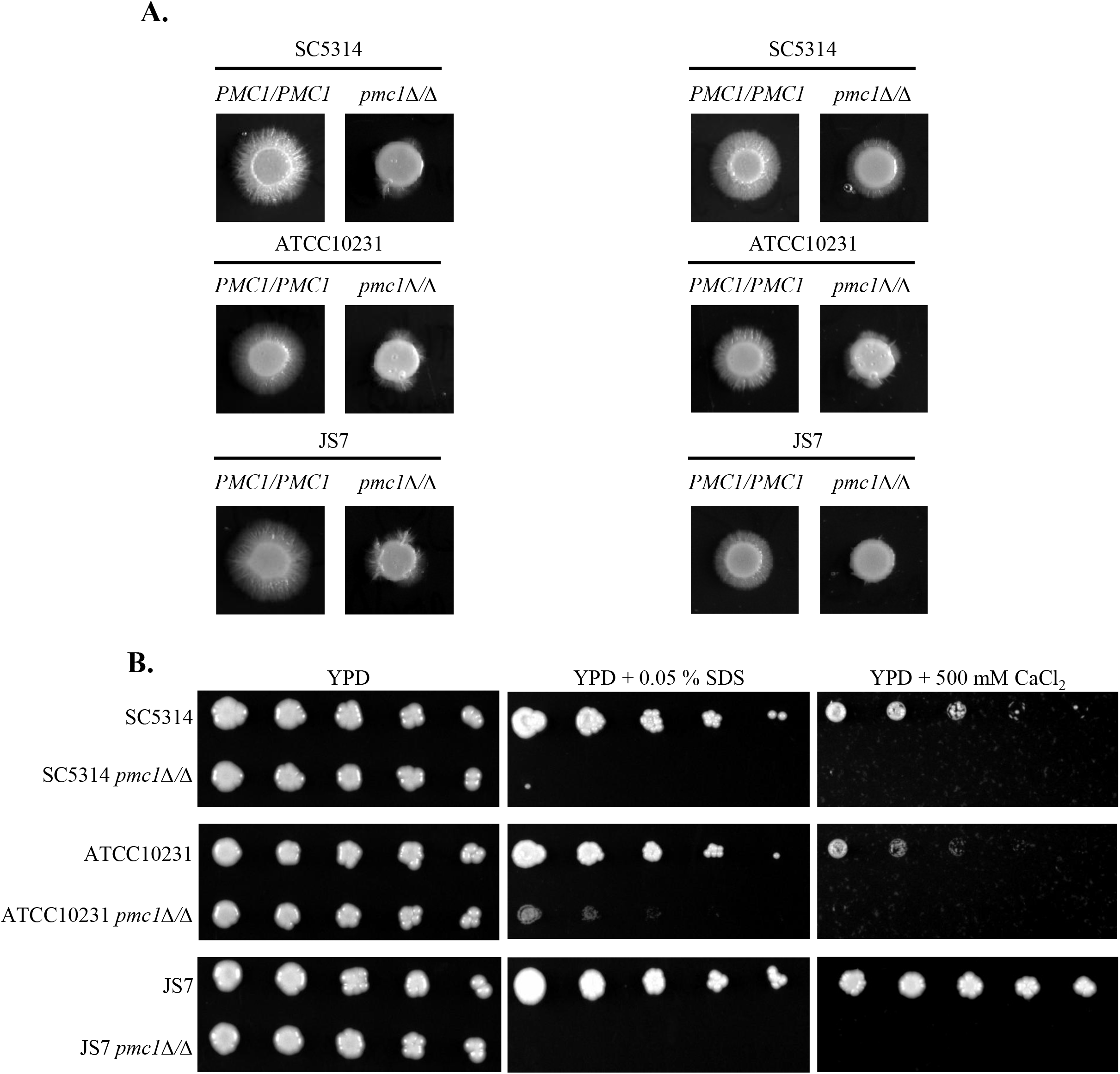
Loss of *PMC1* results in hyphal defects and sensitivities to SDS and calcium. (A) *C. albicans* wild-type and *pmc1Δ/Δ* strains in the SC5314, ATCC10231 or JS7 backgrounds were spotted on agar plates containing 10% FBS (left) or M199 (right). Plates were imaged after 96 hours at 37°C. (B) Spotting assays of indicated strains were performed on YPD agar containing 0.05% SDS or 500 mM CaCl_2_. Plates were imaged after 72 hours at 30°C.

**Supplemental figure 3.**
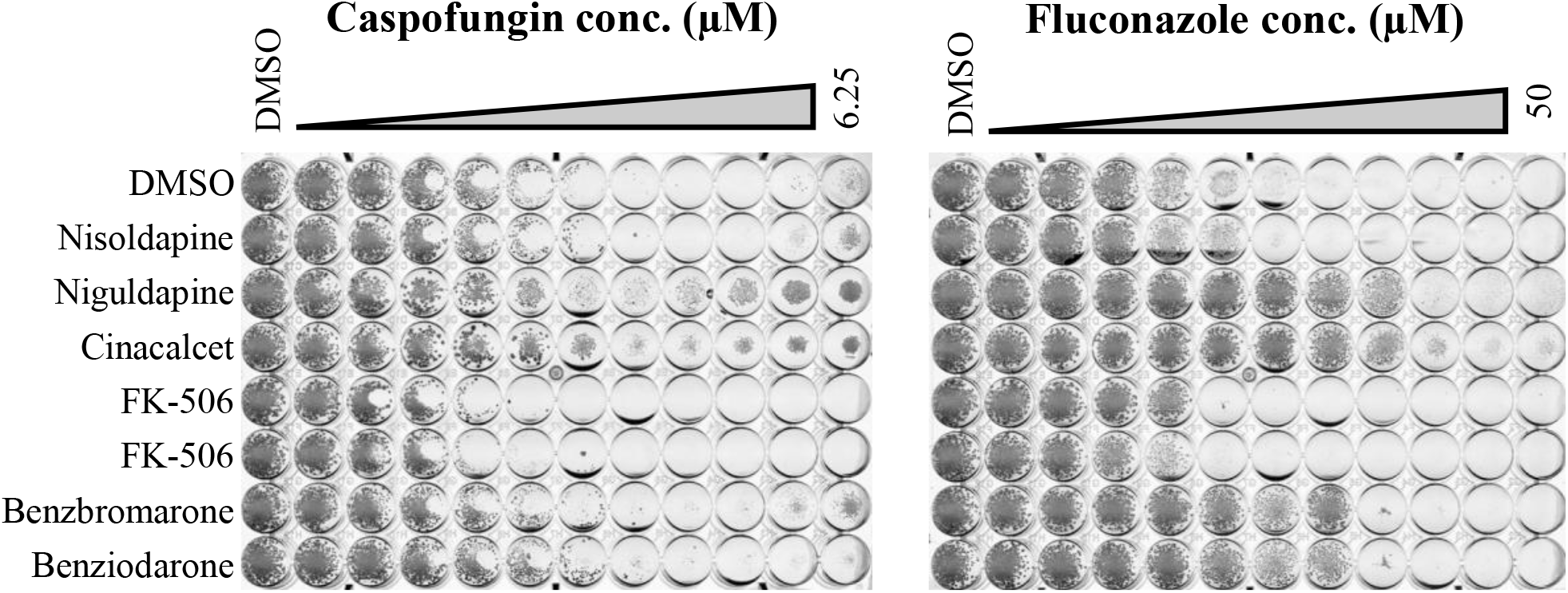
Some drugs targeting calcium homeostasis can affect susceptibility to both azole and echinocandin antifungals. *C. albicans* strain SC5314 was seeded into RPMI medium supplemented with 0.5% DMSO (vehicle control) or 10 µM of the indicated drugs, and dispensed into 96-well plates containing a 2-fold dilution series of caspofungin (left panel) or fluconazole (right panel). Plates were incubated at 35°C and imaged at 48 hours.

